# Temporal interference stimulation targets deep brain regions by modulating neural oscillations

**DOI:** 10.1101/2019.12.25.888412

**Authors:** Zeinab Esmaeilpour, Greg Kronberg, Davide Reato, Lucas C Parra, Marom Bikson

**Affiliations:** Department of Biomedical Engineering, The City College of the City University of New York, New York NY USA; Champalimaud Centre for the Unknown, Neuroscience Program, Lisbon, Portugal

**Keywords:** temporal interference, interferential stimulation, amplitude modulation, gamma oscillation, non-invasive deep brain stimulation

## Abstract

Temporal interference (TI) stimulation of the brain generates amplitude-modulated electric fields oscillating in the kHz range. A validated current-flow model of the human head estimates that amplitude-modulated electric fields are stronger in deep brain regions, while unmodulated electric fields are maximal at the cortical regions. The electric field threshold to modulate carbachol-induced gamma oscillations in rat hippocampal slices was determined for unmodulated 0.05-2 kHz sine waveforms, and 5 Hz amplitude-modulated waveforms with 0.1-2 kHz carrier frequencies. The neuronal effects are replicated with a computational network model to explore the underlying mechanisms. Experiment and model confirm the hypothesis that spatial selectivity of temporal interference stimulation depends on the phasic modulation of neural oscillations only in deep brain regions. This selectivity is governed by network adaption (e.g. GABA_b_) that is faster than the amplitude-modulation frequency. The applied current required depends on the neuronal membrane time-constant (e.g. axons) approaching the kHz carrier frequency of temporal interference stimulation.

## Introduction

Temporal Interference (TI) stimulation delivers high frequency (kHz) sinusoidal stimulation to multiple electrodes on the scalp, where small differences in frequency (e.g. 2 and 2.10 kHz) between electrodes results in an Amplitude-Modulated (AM) electric field deep in the brain (e.g. 2.05 kHz “carrier” whose amplitude is modulated with a “beat” of 100 Hz)^1^. While targeted deep brain structures are exposed to an amplitude-modulated kHz electric fields, superficial cortex is stimulated with higher magnitude unmodulated kHz electric fields. The effectiveness of temporal interference stimulation^2^ thus depends on: 1) steerability of the amplitude-modulated electric fields to targeted deep brain regions^3,4^; 2) the extent to which neuronal activity is more responsive to amplitude-modulated high-frequency electric fields compared to unmodulated electric field (selectivity); 3) the current intensity requirement at the scalp to produce sufficiently strong amplitude-modulated kHz fields deep in the brain (sensitivity).

The effects of electrical stimulation on neuronal oscillations are often analyzed because of their sensitivity to external electric fields^5–7^ and involvement in a broad range of cognitive functions and diseases^8–10^. Conventional transcranial Alternating Current Stimulation (tACS) applies ~2 mA at the scalp, producing electric fields up to ~0.8 V/m in the human brain^11^. In animal models, such small sinusoidal electric fields can modulate oscillations at stimulation frequencies below 100 Hz^6,12–15^ but not evidently for weak kHz frequency stimulation^15,16^. Generally, there is a severe trade-off between the use of kHz stimulation frequencies and amplitudes required for brain stimulation^17,18^. Estimates of the temporal interference electric fields required for acute neuronal modulation in mouse range from 60-350 V/m^1,4^ corresponding to 167-970 mA at the human scalp^19^. Applying kHz tACS with currents of only 1 mA produces mixed effects in human experiments^20^, with loss of efficacy when the waveform is not continuous^21^. The foundation of temporal interference stimulation is the report that neural firing is more sensitive to amplitude-modulated than unmodulated kHz stimulation^1^. However, the low-pass properties of neuronal membranes^15,22^ would a priori predict equal attenuation of both unmodulated kHz and amplitude-modulated kHz stimulation^2,18^ – making amplitude-modulated kHz stimulation as ineffective as unmodulated kHz stimulation. We integrate and reconcile these confounding findings.

Our goal was to develop an experimentally constrained theory for what makes the CNS sensitive to amplitude-modulated high-frequency (kHz) stimulation, how this sensitivity differs compared to unmodulated sinusoidal stimulation at low and high frequencies, and link the sensitivity and selectivity to the spatiotemporal electric fields produced across the brain during temporal interference stimulation. The hippocampal brain slice model is among the most characterized systems in neuroscience and exhaustively tested in screening the effects of electrical stimulation^6,15,23–25^. Specifically, gamma oscillations have been previously shown to be most sensitive to conventional forms of electrical stimulation, with effects reliably predicted by a computational network model^6^. Here, we use this system to test the effects of amplitude-modulated kHz stimulation and contrast outcomes to unmodulated kHz and low-frequency sinusoidal stimulation. We couple this data into a multi-scale model of temporal interference brain current flow and network neuromodulation. We show that temporal interference stimulation depends on the value of phasic modulation of neural oscillations in deep brain regions, as opposed to steady increases driven by unmodulated kHz fields at the cortex. Sensitivity depends on a time constant of membrane polarization close to carrier frequency, while selectivity depends on network homeostatic kinetics that are faster than the frequency (beat) of amplitude modulation.

## Results

### Temporal Interference current flow model

We begin with a computational model of the spatial distribution of electric field across the human brain, using a previously verified modeling pipeline^11,26^. We considered a standard temporal interference montage with two bipolar pairs of electrodes on opposite hemispheres (Figure 1, A.1) applying 1 kHz and 1.005 kHz sinusoidal stimulation with an exemplary amplitude of 167 mA. In regions where electric currents from each electrode pair intersects, the resulting electric field has a carrier frequency of 1.0025 kHz with an amplitude modulation (change in peak electric field) at 5 Hz. At the superficial cortex located near each electrode pair, electric field magnitudes reach peak values of ~80 V/m (Figure 1.A.2). At these locations, the electric field was modulated minimally (amplitude-modulation of ~15%). In contrast, in deep brain regions amplitude-modulation of electric fields could be as high as 50% or more (Figure 1.A.3), corresponding to changes of ~60 V/m at the 5 Hz beat frequency.

**Figure 1:**
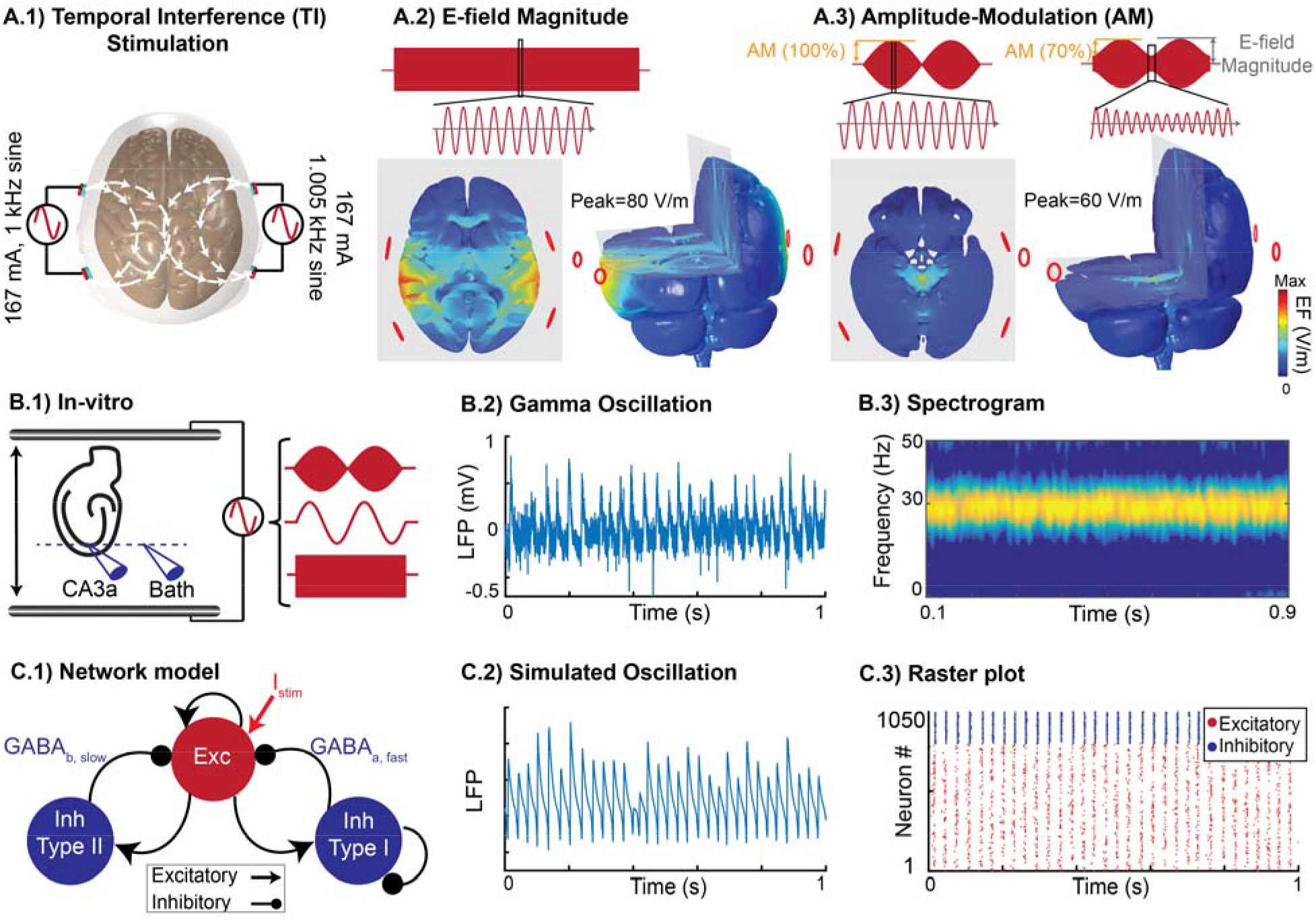
Experimental and computational approaches. **(A)** Computational current-flow model. A.1, Temporal Interference (TI) stimulation via two pairs of electrodes on scalp. Current flows between FT7 and P7 on the left and between FT8 and P8 on the right hemisphere. A.2 Spatial distribution of electric field magnitude in posterior/anterior direction across brain. A.3, Spatial distribution of amplitude-modulation across the brain in posterior/anterior direction. **(B)** Rodent in vitro model of gamma oscillations. B.1, Experimental setup: spatially uniform electric field was applied across hippocampal slice in an interface chamber. Recording of gamma oscillation in CA3a region relative to an isopotential electrode in the bath. B.2 and B.3, Gamma oscillation induced by 20 μM carbachol in vitro and its stability in power and frequency. **(C)** Computational model of gamma oscillations. C.1, The network model has excitatory and inhibitory neurons (1050 neurons, 800 excitatory) that are sparsely connected with varied synaptic weights. C.2, Simulated gamma oscillation in the network model by averaging postsynaptic currents across the network. C.3, Rasterplot representing the firing activity of excitatory (red) and inhibitory (blue) neurons during induced gamma oscillation.

Both the electric field magnitude and amplitude-modulation scale linearly with the applied current. In cortex, unmodulated electric field magnitudes can reach ~0.48 V/m per mA applied current, while in deep brain areas amplitude-modulation of electric fields can reach ~0.36 V/m per mA applied current. Therefore, while amplitude-modulated kHz stimulation can be directed to deep brain regions, on the cortex electric field magnitudes will also be high, consistent with prior models^3,4^.

### Amplitude-modulated and unmodulated kHz stimulation of hippocampal brain slice oscillations

Adapting previous methods ^22^, uniform amplitude-modulated kHz, unmodulated kHz, and low-frequency AC fields were generated across hippocampal slices exhibiting gamma network oscillations (Figure 1.B.1). Consistent with prior reports^6,27^, 20 μM carbachol induced oscillatory activity in local field potentials measured in the CA3a region of the hippocampus (Figure 1.B.2, B.3). Oscillations were typically stable for over 3 hours. Our approach was to systematically contrast the acute effect (2 s, 100 repetitions per slice) of 5 Hz (low), 100 Hz (mid), and 2 kHz (high) frequency sinusoidal unmodulated electric field with 5 Hz amplitude-modulated kHz electric fields with 0.1, 1, or 2 kHz carrier, on gamma oscillations in hippocampal brain slices. For each waveform a range of electric field amplitudes were tested around an empirical threshold range.

We defined two metrics to quantify gamma power: 1) dynamic modulation, which captures fluctuations in gamma power during stimulation; and 2) static modulation, which captures the average gamma power during stimulation (see Methods). Unless otherwise stated, results are reported as mean ± SEM. Low-frequency 5 Hz sine electric field was applied at intensities of 1, 3 and 5 V/m (Figure 2.B.1). There was a monotonic relationship between electric field intensity and dynamic modulation of gamma power, with significant effects for field intensities > 1 V/m (3 V/m: dynamic modulation = 1.36±0.01, n=8; 5 V/m: dynamic modulation=1.65±0.07, n=8) (Figure 2.D.2). The enhancement and suppression during each phase of the 5 Hz stimulation was approximately symmetric, such that there was no significant static modulation (n=8, p=0.5, oneway ANOVA) (Figure 2.D.1, sine 5 Hz).

**Figure 2:**
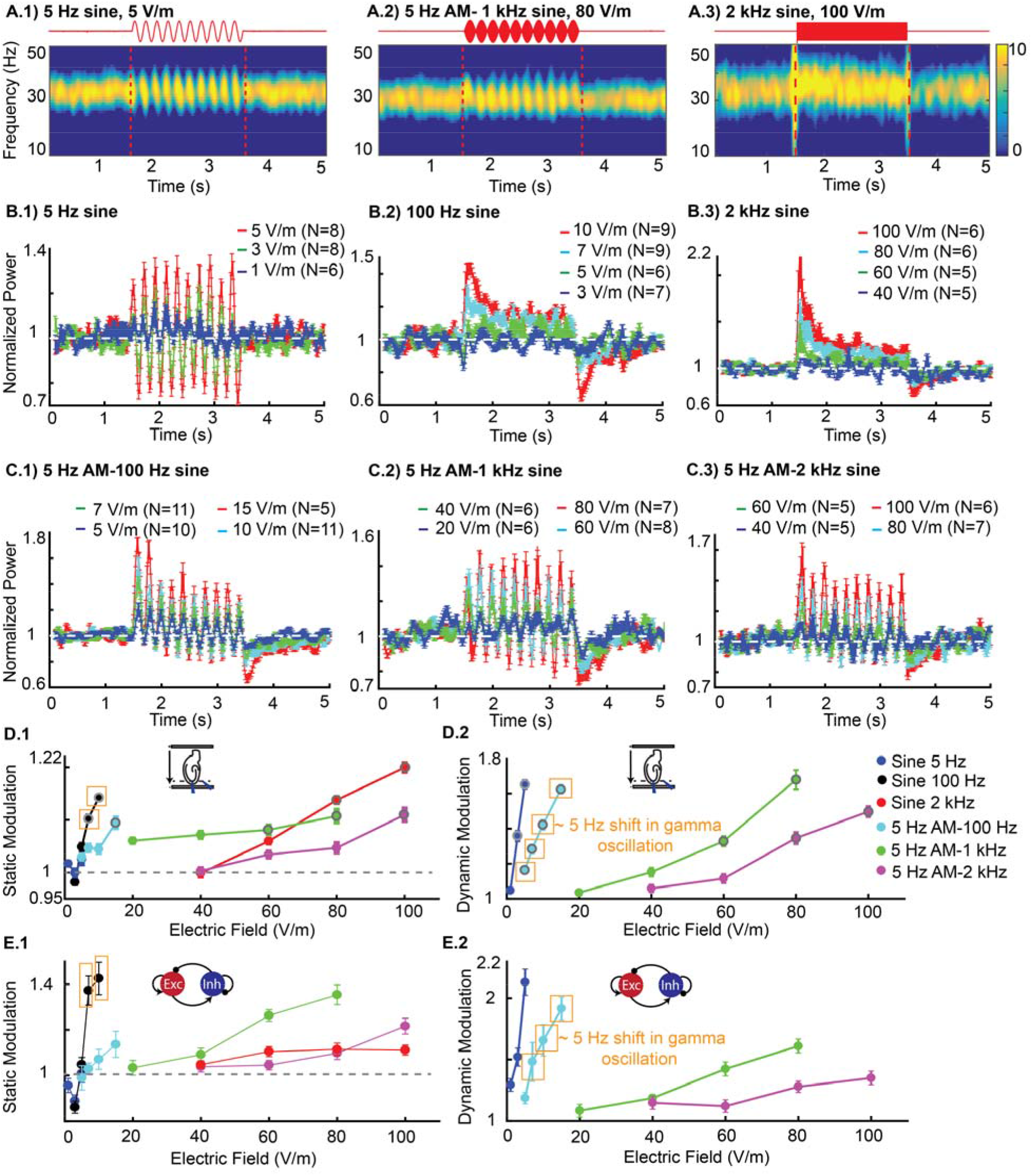
Changes in hippocampal gamma oscillation during application of Amplitude-Modulated high-frequency field as well as low, mid and high-frequency (unmodulated) sinusoidal waveforms in vitro. **(A)** Mean spectrogram of oscillation (in dB) for 2 seconds of stimulation (between 1.5 and 3.5 s) using 5 Hz, 5 V/m sine waveform (A.1), Amplitude-Modulated (AM) waveform, 5 Hz AM-1 kHz sine, 80 V/m (A.2), 2 kHz sine waveform, 100 V/m (A.3). **(B)** Mean (±SEM) of normalized power across slices for different intensities in 5 Hz sine (B.1), 100 Hz sine (B.2), 2 kHz sine (B.3). **(C)** Mean (±SEM) of normalized power across slices for different intensities in amplitude-modulated waveform with 5 Hz envelop and 100 Hz carrier frequency (C.1), 1 kHz carrier frequency (C.2) and 2 kHz carrier frequency (C.3). **(D)** Modulation of gamma power in hippocampal in vitro experiments. D.1) Mean (±SEM) of static modulation of power during stimulation in in vitro experiment measured as mean power modulation during 1 s of stimulation relative to baseline. D.2) Mean (±SEM) of dynamic modulation. In 5 Hz sine stimulation, dynamic modulation calculated as power ratio between interval of positive and negative field and in amplitude-modulated waveforms dynamic modulation is quantified as a ratio of peak (> 50% of peak field intensity) to trough (< 50% of peak field intensity). Error bars indicate standard error of mean. N, number of slices. Grey ring indicates statistically significant modulation relative to baseline, p<0.05; significance was calculated via one-way ANOVA with Tukey post-hoc test. Yellow boxes indicate ~ 5 Hz shift in peak gamma frequency during 100 Hz stimulation (modulated and unmodulated)**. (E)** Modulation of power during stimulation in computational network model.

Stimulation with unmodulated sinusoids at mid (100 Hz) and high frequencies (2 kHz) did not produce significant changes in hippocampal gamma oscillations using intensities effective for low-frequency stimulation (i.e. ≤ 5 V/m). Significant steady increases (static modulation) in gamma power were detected using electric field intensities ≥ 7 V/m for 100 Hz (7 V/m: static modulation=1.11±0.06, n=9, 10 V/m: static modulation=1.14±0.037, n=9) (Figure 2 B.2) and electric field intensities ≥ 80 V/m for 2 kHz (80 V/m: static modulation=1.15±0.07, n=6, 100 V/m: static modulation=1.20±0.01, n=6) (Figure 2 B.3). As tested, 2 kHz stimulation did not change the frequency of peak gamma power, while 100 Hz stimulation shifted ongoing oscillation frequency by ~5 Hz. Static modulation increased with increasing electric field intensity (Figure 2.D.1) and decreased with increasing stimulation frequency. Using higher frequencies, stronger field intensities are required to produce the same effect (Figure 2.D.1). Stimulation with unmodulated sinusoids at frequencies of 100 Hz and 2 kHz did not produce dynamic modulation of oscillations-this result is expected since these waveforms include no low-frequency (e.g. 5 Hz) amplitude modulation.

Stimulation with 5 Hz amplitude-modulated waveforms resulted in dynamic modulation of hippocampal gamma activity at the 5 Hz “beating” frequency (Figure 2.A.2). The magnitude of this dynamic modulation of oscillations increased with electric field magnitude and decreased with carrier frequency (Figure 2.D.2). 5 Hz amplitude-modulated stimulation produced significant dynamic modulation using electric field intensities ≥ 5 V/m for 100 Hz carrier (5 V/m: dynamic modulation=1.16±0.08, n=10; 7 V/m: dynamic modulation =1.280.12, n=11; 10 V/m: dynamic modulation =1.42±0.14, n=11; 15 V/m: dynamic modulation=1.61±0.06, n=5). Electric fields greater than ≥ 60 V/m were effective with a 1 kHz carrier (60 V/m: dynamic modulation=1.33±0.10, n=8; 80 V/m: dynamic modulation=1.68±0.03, n=7). Electric fields ≥ 80 V/m were effective with a 2 kHz carrier (80 V/m: dynamic modulation=1.35±0.11, n=7; 100 V/m: dynamic modulation =1.50±0.14, n=6). Amplitude-modulated stimulation with 1 kHz and 2 kHz stimulation did not change the frequency of peak gamma power, while 100 Hz carrier stimulation shifted ongoing oscillation frequency by ~5 Hz.

Stimulation with 5 Hz amplitude-modulated waveforms also produced significant static modulation for intensities ≥ 15 V/m for 100 Hz carrier (15 V/m: static modulation=1.10±0.06, n=5), ≥ 60 V/m for 1 kHz carrier (60 V/m: static modulation=1.09±0.09, n=8; 80 V/m: static modulation=1.11±0.09, n=7) and ≥ 100 V/m for a 2 kHz carrier (100 V/m: static modulation=1.10±0.01, n=6); showing a non-symmetric effect on gamma power modulation. Hippocampal gamma oscillation sensitivity to amplitude-modulated waveforms decreases with increasing carrier frequency; so stronger stimulation intensities are required to modulate gamma activity when higher carrier frequencies are used (Figure 2.D.2).

### Computational network model of hippocampal gamma oscillation

It is well known that the sensitivity of transmembrane potentials to sinusoidal electric fields decreases with increasing stimulation frequency^15^ which is explained by the membrane time constant^15,22^. In active networks, sensitivity to electric fields is further dependent on network dynamics^6,15,28^. But the implications of these prior findings to amplitude-modulated kHz electric field has remained unclear. We adapted a previously verified computational network model of hippocampal gamma oscillations. The model uses single-compartment excitatory and inhibitory neurons, which are coupled to the external electric field^6^. The computational model provides quantitative predictions for the sensitivity of gamma oscillations to unmodulated and amplitude-modulated stimulation across frequencies. Two key modifications to the prior model were implemented: 1) in addition to fast synaptic inhibition, motivated by typical *GABA_a_* receptors ^6,29^, we also included a slower *GABA_b_*-type inhibitory conductance with higher activation threshold ^30,31^; 2) the membrane time constant (*τ*) was decreased to 1 ms. We go on to show that these properties are essential to sensitivity and selectivity of temporal interference stimulation.

For low-frequency 5 Hz sine, the model reproduced experimental modulation of hippocampal gamma power (Figure 2.E1, E2, sine) as shown previously^6^. For higher frequency stimulation (both amplitude-modulated and unmodulated), the model captures major features of our in vitro experiments: 1) the inverse relationship between stimulation carrier frequency and the sensitivity of gamma oscillations to stimulation (i.e. much higher electric field magnitude required for high frequency stimulation (Figure 2.E1, E2)); 2) for a given carrier frequency, static modulation and dynamic modulation increased with field magnitude (Figure 2.E.1, E2); 3) stimulation with the 100 Hz carrier shifted gamma oscillation frequency, while 1 and 2 kHz carriers did not produce significant change in gamma frequency.

For stimulation with unmodulated mid (100 Hz) and high (2 kHz) frequency electric field, the model also reproduced the specific time course of gamma power modulation in our experiments. There is a rise in gamma power at the onset of stimulation, followed by a decay to steady state, which remains above baseline (Figure 3.B). We only observe increases in gamma power, reflecting the sensitivity of the active network to the depolarizing phase of the sinusoidal electric field waveform. Indeed, the response profile in unmodulated high frequency stimulation is similar to DC depolarizing stimulation^6^. The observed time constant of network adaptation is governed by recruitment of high-threshold inhibitory neurons, which produce slow *GABA_b_* post-synaptic inhibition^30^(Figure 3.A3).

**Figure 3:**
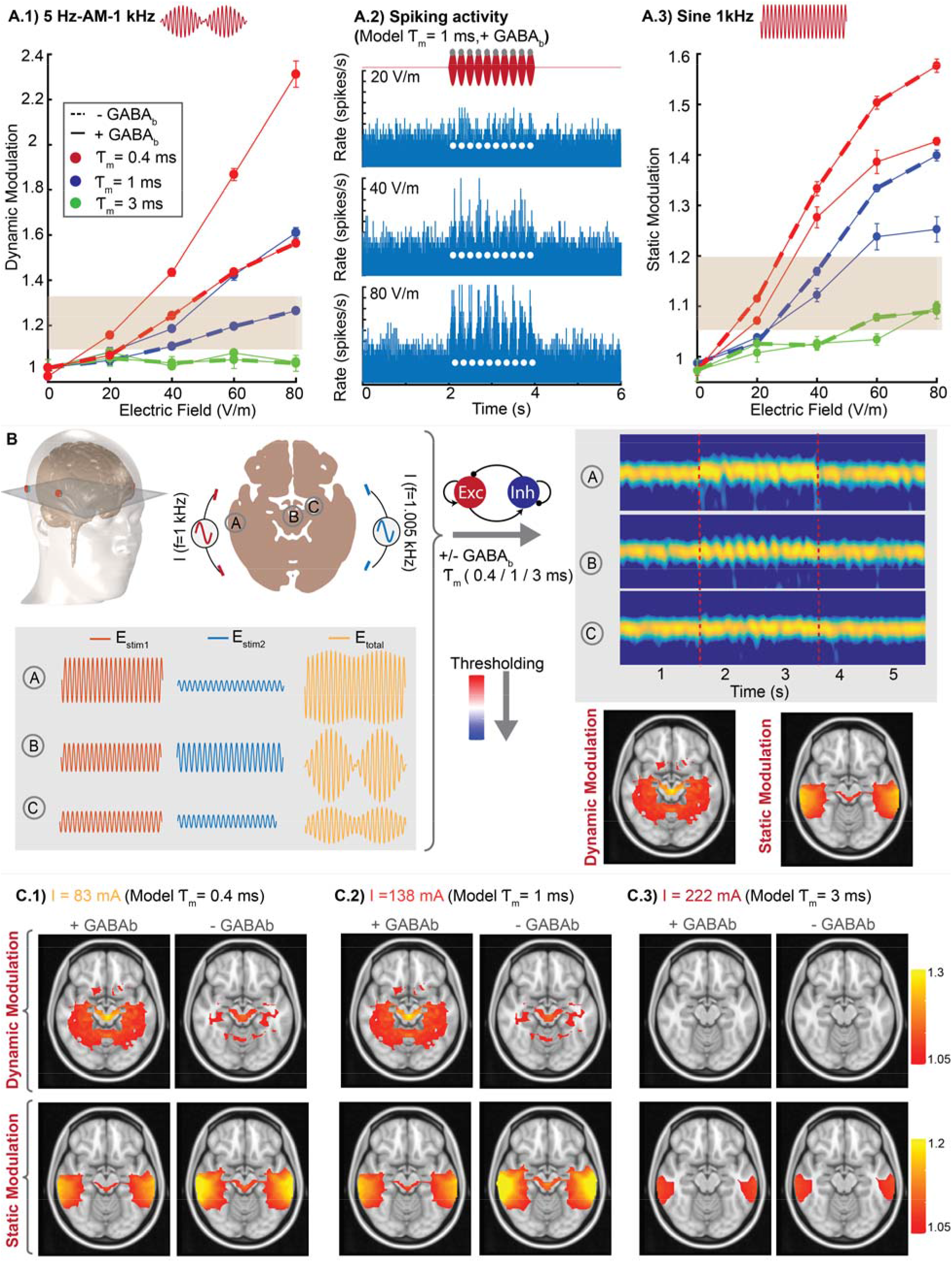
Generalized network model of temporal interference stimulation of oscillation. (A.1) Effect of membrane time constant (*τ_m_*) and GABAergic inhibition on gamma modulation using amplitude-modulation (AM) waveform (5 Hz envelop and 1 kHz carrier frequency) with different field intensities. Each point represents mean (±SEM) of normalized gamma power for repeated runs of model (Blue solid line matches the experimental data of hippocampal gamma). A.2) Spiking activity for 5 Hz-AM-1 kHz stimulation using different electric field intensities. Y axis is clipped for illustration purposes in 80 V/m. A.3) Effect of membrane time constant and GABA_b_ on gamma modulation using unmodulated 1 kHz sine waveform with different electric field intensities. (**B)** Workflow for multi-scale model of dynamic and static modulation of oscillations across brain. Electric field for each voxel of brain was calculated using computational head model and used as I_stim_ in network model to generate corresponding modulation (both static and dynamic). (**C)** Model predictions for dynamic and static modulation using different network model parameters: *τ_m_*= 0.4 ms with/without GABA_b_ inhibition (C.1), *τ_m_*= 1 ms with/whiteout GABA_b_ inhibition (C.2), *τ_m_*= 3 ms with/without GABA_b_ inhibition (C.3). Grey box in A.1 and A.3 indicates thresholds derived from experimental hippocampal recordings used in plotting static and dynamic modulation in panel C. White circles in A.2 indicates peaks of amplitude-modulated waveform.

Stimulation of 5Hz-amplitude-modulated waveforms modulated ongoing hippocampal gamma oscillation at the envelop frequency (5 Hz) as observed in vitro. Notably dynamic modulation due to amplitude-modulated waveforms is greater than the static modulation due to the corresponding unmodulated waveform. In the model this difference depends on the presence of *GABA_b_* synapses, which control the timescale of network adaptation (Figure 3.A1).

### Generalized model of network oscillation sensitivity and selectivity to Temporal Interference stimulation

The above model was built to match experimental data in hippocampal CA3 slices. To generalize the model, we consider how its predictions depend on details of the model’s biophysical parameters. We specifically considered changes in membrane time constant and GABAergic inhibition (GABA_b_), with alterations described relative to the computational network parameters that reproduced hippocampal gamma oscillations (*τ_m_* = 1 *ms*, + *GABA_b_*, solid blue line).

Decreasing the membrane time constant increases the sensitivity of gamma oscillations to both amplitude-modulated and unmodulated kHz electric fields (Figure 3.A.1, 3.A.3; solid red line, *τ_m_* = 0.4 *ms*). Conversely, increasing the membrane time constant leads to insensitivity of gamma oscillations to stimulation waveforms in the electric field range tested in experiments (Figure 3.A, *τ_m_* = 3 *ms*, solid green line). In our model, the single-compartment membrane time constant reflects the most sensitive neuronal element to kHz stimulation. A membrane time constant of 1 ms reproduces our experimental data which is moderately faster than prior simulations of gamma oscillations (*τ_m_* = 3-10 ms^29,32,33^) and the resting state somatic polarization time constant by electric fields (*τ_m_*~20 ms^6,15^). This finding has direct implications for the neuronal element targeted by interferential stimulation, namely axons^25,34^.

Spiking activity in the model suggests a slight increase in unit activity even at an electric field intensity below the thresholds for modulating network gamma oscillations (figure 3.A.2). It is expected (the most sensitive) individual neurons will respond to electric field before changes in network power exceed a threshold^6^.

Removing GABA_b_-mediated inhibition decreases dynamic modulation in response to amplitude-modulated waveforms, while increasing static modulation in responses to unmodulated sinusoidal electric fields (compare solid lines with dashed lines, Figure 3.A). The GABA_b_ synapses are therefore critical for the network to exhibit sensitivity and selectivity for amplitude-modulated waveforms.

Finally, we determined how the sensitivity and selectivity to temporal interference stimulation across the brain are governed by cellular and network biophysics (figure 3.B). To do so we assume a generic neural circuit, which reoccurs throughout the brain (i.e. at each voxel), simulated with the experimentally validated model described above. Of course, not all brain regions are identical, but such an approach allows a principled analysis of the parameters governing sensitivity/selectivity of the brain to temporal interference stimulation.

The electric fields generated in each brain region (2×2×1 mm voxel) of the temporal interference current flow model was used as the input (I_stim_) in the network model of gamma oscillations. The electric field varied in both peak intensity and the degree of amplitude-modulation across regions, which produces a mix of static and dynamic neuromodulation at each brain region (e.g. exemplary regions A, B, C). Mapping of static and dynamic modulation in each region is then represented using thresholds derived from experimental hippocampal recordings. In this series only the cellular and network biophysics are varied, with the applied stimulation current (I) selected to demonstrate sensitivity and selectivity (figure 3 C).

To test the robustness of the multiscale model and to identify which features govern sensitivity and selectivity, we varied parameters that were found to be critical for determining static and dynamic neural modulation in the network model (i.e. *τ_m_*, *GABA_b_*). A faster membrane time constant reduces the minimum stimulation current required to modulate oscillations in deep regions (i.e. increases sensitivity to temporal interference stimulation; figure 3C, compare panels 1,2,3). However, even for the most sensitive parameter choice (figure 3.C1, *τ_m_* = 0.4 *ms*), this threshold stimulation current is still much higher than conventional tACS (i.e. 83 mA vs. 2 mA).

Removing GABA_b_ inhibition reduced hot spots of dynamic modulation in (target) deep brain regions, while increasing static modulation in overlying cortex (figure 3C.1–3C.3 compare left/right panels). Therefore, GABA_b_ inhibition improves selectivity for deep brain regions. However, the model predicts that dynamic modulation in deep brain regions (figure 3.C, top row) is generally associated with static modulation of cortical areas (figure 3.C, bottom row). This indicates that selectivity is rather limited regardless of parameter choices.

## Discussion

Temporal interference stimulation has been promoted as a tool to selectively modulate neural activity in deep brain regions^1^. The ability of temporal interference stimulation to achieve such selectivity depends on 1) the relative magnitude of amplitude-modulated kHz electric fields (in deep brain) as compared to the unmodulated kHz electric fields (in cortex), and 2) the sensitivity of regional neural networks to amplitude-modulated kHz electric field in contrast to unmodulated kHz waveforms. With regards to the electric field magnitude during temporal interference stimulation, this can be predicted with validated finite element head models (Figure 1.A). With regards to network sensitivity, here we calibrate the dose-response to amplitude-modulated kHz and unmodulated kHz waveforms in a canonical model of hippocampal gamma oscillations (Figure 2). We show how cellular and network biophysics, namely the time constants of axonal membranes and GABAergic inhibition, can explain (Figure 2) and generalize (Figure 3) both sensitivity and selectivity to temporal interference stimulation. Integrating experimentally-verified current flow and network models, we can make predictions about the ability of temporal interference stimulation to selectively target deep brain regions.

Starting with hippocampal gamma oscillations, we show an amplitude-modulated kHz (1 kHz modulated at 5 Hz) electric fields of ~60 V/m is required for significant dynamic modulation of oscillations (Figure 2.D.2), while an unmodulated kHz electric field of the same intensity (~60 V/m) is required to produce static modulation (Figure 3.A3). Assuming these biophysics are uniform across the brain, selectivity therefore requires that the amplitude-modulated kHz electric field magnitude in deep brain regions be greater than ~60 V/m, while the unmodulated kHz electric field in the cortex is less than ~60 V/m. Is this achievable with temporal interference stimulation? With a basic dual bipolar electrode configuration, producing 60 V/m in deep brain regions requires ~167 mA on the scalp (Figure 1.A.3), corresponding to a peak unmodulated electric field of ~80 V/m at the cortex (Figure 1.A.2). Making it unlikely for temporal interference stimulation to produce dynamic modulation with amplitude-modulated kHz electric field in deep brain regions without also producing static modulation of the overlying cortex with unmodulated kHz electric fields. This prediction holds across a range of neuronal and network biophysics, under the assumption that they are uniform across the brain (Figure 3.C). However, selective deep brain stimulation by temporal interference stimulation derives from: 1) use of more sophisticated electrode montages^3,19^; 2) cellular or network features special to deep brain regions; or 3) impact (value) of dynamic oscillations in deep brain regions versus static modulation at superficial cortex. Ascendant to any approach to temporal interference stimulation, we show that the sensitivity (applied current required) and selectivity (responsiveness to amplitude-modulated verse unmodulated electric) of the brain to temporal interference stimulation is governed by neural-compartment and network-oscillation features identified here.

What explains the sensitivity of the brain to temporal interference? Amplitude-modulated kHz stimulation has frequency content around the carrier frequency, not at the beat frequency, such that the low-pass filter properties of neuronal membranes^15,22^ will attenuate amplitude-modulated kHz similarly to unmodulated kHz^2,18^. Whereas prior neuron models of low-frequency stimulation (tDCS, tACS) considered polarization of somatic ^6,18^ or dendritic compartments^24,35–37^, here we consider axonal polarization. Axons not only have the highest sensitivity to stimulation (polarization coupling constant 4x of somas^34^) they also have the fastest time constants. A membrane time constant not exceeding 1 ms is pivotal to sensitivity to kHz carriers (Figure 3.A) implicating axons as the temporal interference stimulated neuronal element. Active networks provide additional amplification by effectively increasing the polarization coupling constant^6,38^, and through non-linear network responses^12,13,39,40^. Characterizing what determines the sensitivity of deep brain regions to amplitude-modulated kHz stimulation should consider how axons are polarized^41,42^ and network amplification factors.

With regard to selectivity, we note even conventional tES easily reaches deep brain structures^11,14,43,44^ with some deep selectivity achievable with High-Definition (HD) optimization^3,19^. Temporal interference stimulation offers possibilities to further improve selectivity, but this is subject to constraints on current flow patterns^3,4^ (Figure 1.A) and the potency of amplitude-modulated electric fields compared to unmodulated electric fields^2^. Here this potency is largely determined by the magnitude and time constant of a network homeostatic adaptation mechanism. For unmodulated kHz electric fields (e.g. in cortex), this adaptation *suppresses* the degree of static modulation. For amplitude-modulated kHz electric fields (e.g. in deep brain), this adaptation *boosts* the degree of dynamic modulation. Here we attribute this adaptation mechanism to GABA_b_ synapses^30^, though other cellular and network adaptation mechanisms exist^45^. Only adaptation on a timescale faster that the amplitude-modulation “beat” frequency should enhance dynamic modulation and selectivity. Identifying brain regions, neurons, or parallel interventions that facilitate this adaptation may improve temporal interference effectiveness.

Cellular and networks biophysics are not uniform across the brain, and moreover would change with brain state^46^ and disease^47^. Similarly, divergent results across animal studies (e.g. high selectivity of amplitude-modulated kHz^1^; or low sensitivity to tACS^48^) may be explained by variability in these governing parameters. With high-intensity electric fields alternative biophysics such as ion accumulation^49^, fiber block^50–52^, asynchronous firing^53^ and/or heating^54^ may also be considered, but were not required to explain our results. Electrophysiology experiments with kHz electric field stimulation must carefully account for the fidelity of delivered stimulation and recording artifacts^55^. Indeed, we avoided intracellular electrodes because microelectrodes can collect current from kHz stimulation^56^ risking either artifactual intracellular stimulation (as the microelectrodes acts as a collector “antenna”) or amplifier distortion^57^.

We show selective stimulation of deep brain regions derives from phasic modulation of neuronal oscillation with dynamics adapting faster than the “beat” frequency of temporal interference stimulation. An outstanding question for effectiveness is what is the required stimulation intensity? Oscillations provide some amplification, but sensitivity is ultimately throttled by membrane constant (e.g. > 80 mA current for 0.4 ms time constant; Figure 3.C). Effective temporal interference with intensities comparable to conventional tACS (~2 mA) would require still faster effective membrane time constants based on mechanisms that remain speculative for low-intensity amplitude-modulated kHz fields such as polarization of intracellular compartments, transverse axonal polarization^58^, or non-neuronal cellular targets^59^. Alternatively, meaningful changes in cognition could derive through modulating a small fraction of hyper-responsive neurons which are part of oscillating networks. Notwithstanding technical and conceptual limitations, we show the cellular mechanisms of temporal interference stimulation of the brain can be addressed through the sensitivity and selectivity of neuronal oscillations.

## Materials and Methods

All animal experiments were carried out in accordance with guidelines and protocols approved by the Institutional Animal Care and Use Committee at The City College of New York, CUNY.

### Hippocampal slice preparation

Hippocampal brain slices were prepared from male Wistar rats aged 3–5 weeks old, which were deeply anaesthetized with ketamine (7.4 mg kg-1) and xylazine (0.7 mg kg-1) and sacrificed by cervical dislocation. The brain was quickly removed and immersed in chilled (2–6 °C) dissecting solution containing (in mM) 110 choline chloride, 3.2 KCl, 1.25 NaH2PO4, 26 NaHCO3, 0.5 CaCl2, 7 MgCl2, 2 sodium ascorbate, 3 sodium pyruvate, 10 D-glucose. Transverse hippocampal slices (400 μm thick) were cut using a vibrating microtome (Campden Instruments, Leicester, England) and transferred to a recovery chamber for 30 minutes at 34 °C with a modified artificial cerebrospinal fluid (ACSF) containing (in mM) 124 NaCl, 3.2 KCl, 1.25 NaH2PO4, 26 NaHCO3, 2.5 CaCl2, 1.3 MgCl2, 2 sodium ascorbate, 3 sodium pyruvate, and 25 D-glucose. Slices were then transferred to a holding chamber for at least 30 minutes (or until needed) at 30 °C with ACSF containing (in mM) 124 NaCl, 3.2 KCl, 1.25 NaH2PO4, 26 NaHCO3, 2.5 CaCl2, 1.3 MgCl2, and 25 d-glucose. After 60 min, slices were then transferred to a fluid–gas interface recording chamber (Hass top model, Harvard Apparatus, Holliston MA, USA) at 34 °C. All solutions were saturated with a gas mixture of 95% O2–5% CO_2_. All reagents were purchased from Sigma Aldrich (St. Louis MO, USA). Gamma oscillations were induced by perfusing the slices with ACSF containing 20 μM carbachol (carbamoylcholine chloride).

### Extracellular recordings

Recordings of extracellular field potentials in the pyramidal layer of CA3a region of hippocampus were obtained using glass micropipettes (15 MΩ pulled on a P-97, Sutter instruments) field with ACSF. Data acquisition and electrical stimulation were controlled by Power1401-625 kHz hardware and Signal software Version 6.0 (Cambridge Electronic Design (CED), Cambridge, UK). Voltage signals were amplified (10x), analog low pass filtered (20 kHz; Model 3000 differential amplifier, A-M systems, Carlsberg WA, USA) and digitized (20 kHz, Power1401-625 kHz and Signal, CED, Cambridge, UK). To reduce noise and stimulation artifacts, the voltage recordings were always performed relative to an iso-potential electrode placed in bath (Figure 1, B.1). Field recordings overcome potential limitations of intracellular recording under kHz field such as current collection by the capacitive-walled microelectrode leading to artifactual intracellular stimulation ^56^ or possible amplifier distortion^57^.

### Electrical field stimulation

Under the quasi-uniform assumption^60^, the electric field amplitude and waveform generated in a brain ROI can be applied across an in vitro system. Spatially uniform electric fields were applied to slices with varying frequencies and intensities by passing current between two parallel Ag–AgCl wires (1 mm diameter, 12 mm length, 10 mm apart) placed in the recording chamber on opposite sides of the brain slice^6,22^. Field waveforms were generated by function generator (Arbitrary function generator, AFG1062, 60MHz, 300Ms/s, Tektronix, USA) and converted to a controlled current source stimulus by a custom high bandwidth voltage-controlled isolated current source^56^. Unless otherwise stated, the electric field reported throughout the manuscript is the peak electric field for each waveform. Slices were oriented so that the resulting electric field was parallel to the main somato-dendritic axis of CA3a pyramidal neurons (perpendicular to pyramidal cell layer). Before each recording, the applied current intensity was calibrated by measuring the electric field (voltage difference between two recording electrodes separated by 0.4 mm in the slice)^61,62^. Stimulation was applied 30-45 min after application of carbachol when the intensity and frequency of oscillations were stabilized.

### Power analysis and statistics

Signals were recorded in frames of 7 s (1.5 s before and 3.5 s after stimulation) and stimulation was applied for 2 s. Stimulation artifacts were minimized by subtracting the voltage in an iso-potential refence electrode from the recording electrode in the slice (Figure 1). Spectrograms were computed (200 ms hamming window, 90% overlap) on individual 7 s frames and averaged over 100 frames for each stimulation condition (i.e. frequency, waveform and amplitude). Normalized power was measured as a power ratio normalized by prestimulation power in the frequency band of the endogenous oscillation. In case of 100 Hz stimulation (i.e. sine 100 Hz and 5 Hz-AM-100 Hz), which caused ~ 5 Hz shift in endogenous gamma oscillation during stimulation, gamma power was measured and quantified in the center frequency of the oscillation specific to each interval (i.e. baseline, stimulation).

To quantify the mean effect of stimulation we defined the static modulation as the mean power in the gamma band (20-40 Hz) during the final 1 s of stimulation in each frame divided by the mean gamma power immediately preceding stimulation (1 s). To capture the dependence of gamma modulation on the phase of the stimulation waveform, we defined a dynamic modulation metric. For 5 Hz sine stimulation, dynamic modulation was the power ratio of positive field over negative fields. For AM-high-frequency stimulation, dynamic modulation was the ratio of the gamma power during the peak interval to gamma power during the trough interval. Unless otherwise stated, results are reported as mean ± SEM; n= number of slices. Significance (p < 0.05) was characterized by one-way ANOVA followed by post hoc test with Tukey for multiple comparisons. All statistical analysis is done in R.

### Computational head model

We adapted an existing detailed head model with 1 mm resolution to predict the spatial distribution of electric fields across the human brain during temporal interferential stimulation. Briefly, the model was segmented into tissues with different conductivity (scalp, fat, skull, CSF, air, grey and with matter). The model was meshed using ScanIP and solved using a finite element modeling software (COMSOL). We used two independent pairs of electrodes: FT7 and P7 on the left side and FT8 and P8 on the right hemisphere. The spatial distribution of amplitude-modulated electric field was measured in the posterior/anterior direction (see ^63^ for technical details).

### Network model

A network of excitatory and inhibitory neurons was used to explain our results in hippocampal brain slices. The local recurrent CA3 circuit was simulated using a model consisting of 800 excitatory and 250 inhibitory neurons (200 form synapses with *GABA_a_* dynamic, 50 from synapses with *GABA_b_* dynamics). Each cell was modeled as a single-compartment, adaptive exponential integrate-and-fire neuron (AdEx) since it can produce a large variety of neuronal behaviors by changing few parameters ^64^. The following differential equations describe the evolution of the membrane potential V(t) of each neuron:

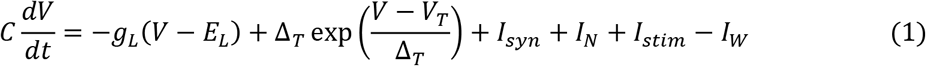

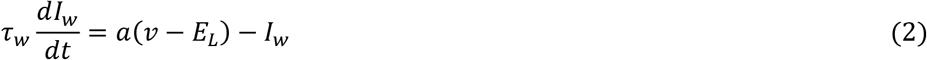

When the current drives the potential beyond *V_T_*, the exponential term actuates a positive feedback which leads to upswings of the action potential. The upswing is stopped at a reset threshold which we fix at *V_thre_* = −50 *mV* and membrane potential is replaced by *V_reset_* and *I_w_* is increment by an amount *b* on the following step:

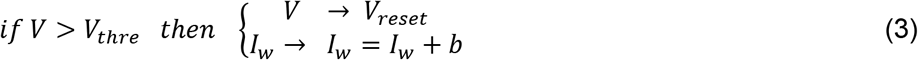

We considered a regular spiking neuron for excitatory cells and a fast spiking neuron model for inhibitory. Parameters are as follows: *Excitatory neurons*: *C* = 100 *pF*, *E_L_* = −55 *mV*, *a* = 2, *b* = 8, *τ_w_* = 400, Δ*T* = 2.7, *g_L_* = 100 *nS*, *V_T_* = −52 *mV*, *V_reset_* = −55 *mV*; *Inhibitory neurons*: *C* = 100 *pF*, *E_L_*(*GABA_a_*) = −62 *mV*, *E_L_*(*GABA_b_*) = −67 *mV*, *a* = 0, *b* = 0, *τ_w_* = 400, Δ_*T*_=1, *V_T_* = − 55 *mV*, *V_reset_*(*GABA_a_*) = −62 *mV*, *V_reset_*(*GABA_b_*) = −67 *mV*

The network was structured such that neurons were connected randomly with uniform probability *p_ij_* of connection between a postsynaptic neuron *i* and a presynaptic *j*, which depended on the type of pre and post-synaptic neuron: *p_EE_* = 0.15, *p_I_GABA a_,E_* = 0.4, *p_I_GABA b_,E_*=1, *p_E,I_GABA a__* = 0.4, *p_E,I_GABA b__* = 0.4, *p_I_GABA a_,I_GABA a__* = 0.4, *p_I_GABA b_,I_GABA a__* = 0, where *E* represents excitatoy and *I* represents Inhibitory. The connectivity was sparser between excitatory neurons than other pairs^28,65^. The synaptic current *I_syn,i_* received by neuron *i* is the result of spiking activity of all connected pre-synaptic neurons *j* which can be decomposed into excitatory and inhibitory components: 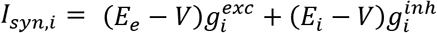. We modeled *g* as decaying exponential function that takes the kicks in at each spike firing of presynaptic spike:

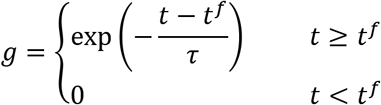

The total inhibitory and excitatory conductance that neuron *i* receives calculated as follows:

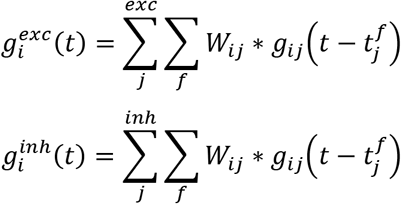

Where *τ_exc_* = 5 *ms*, *τ_inh_GABAa__* = 8 *ms*^32^ and *GABA_b_* conductance has 50 ms of delay and a longer time constant (*τ_inh_GABAb__* = 50*ms*)^66^. The synaptic strengths were chosen to be uniformly distributed for *w_E,E_* ∈ [0,0.3], *w_i_GABAa_,E_* ∈ [0,2], *w_E,i_GABAa__* ∈ [0.5,2.5], *w_i_GABAa_i_GABAa__* ∈ [0,0.5], *w_E,i_GABAb__* ∈ [0,0.5], *w_i_GABAb,E__* = 0.5. In the absence of synaptic input from the network, each excitatory cell is subjected to Gaussian noise (SD=0.5 nA) to simulate spontaneous activity of pyramidal cells under carbachol perfusion (*I_N_*). local field potential (LFP) is thought to result from synaptic activity and we modeled LFP signal by averaging all excitatory and inhibitory postsynaptic currents from the network^6^.

### Model of Electric field in the network

The effect of stimulation was implemented as a small current (*I_stim_*) injected into excitatory neurons^6,18,67^. This approach captures the induced membrane polarization of the single compartment due to external electric field application. Inhibitory neurons were not polarized by the field, assuming a typical symmetric morphology^68^. It has been shown that 1 V/m produces 0.2 mV polarization in low frequency (<7 Hz)^15^. In the model we assumed the DC current amplitude that produces 0.2 mV membrane polarization is equivalent to a 1 V/m electric field (*I*_0.2 *mV polarization in DC*_ = 1 *V*/*m*). The waveform of AM high-frequency stimulation was constructed by subtracting two sinusoidal waveforms where *f_c_* is the carrier frequency and *f_m_* is the modulating frequency (*f_m_* = 5 *Hz*, *f_c_* = 0.1,1,2 *kHz*).
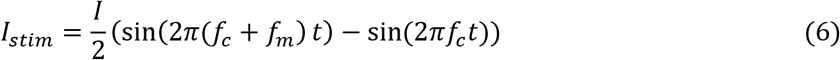

### Generalized model

For all the conditions in the generalized model, network structure (connectivity and synaptic weights) followed the same probability distributions as described above in the network model that represented in vitro experiments. In order to evaluate the effect of membrane time constant on network sensitivity to temporal interreference stimulation, membrane capacitance (C) was changed only for excitatory cells, since inhibitory neurons do not get polarized during stimulation. In the most sensitive network, membrane capacitance was modeled as *C* = 40 *pF* whereas in the less sensitive network membrane capacitance was assumed *C* = 300 *pF*. For studying selectivity, we removed GABA_b_ inhibition by setting the weight of all GABA_b_ synapses to zero (*w_E,i_GABAb__*, = 0). When varying parameters in the model (C, GABA_b_), the noise current simulating the effect of carbochol in pyramidal cells (*I_N_*) was adjusted to keep firing the rates of excitatory and inhibitory cells and the network oscillation frequency within the range of reported experimental data^27,69^. In figure 3, we set a threshold level of static and dynamic modulation at 5 %. This was the maximum amount of modulation that still could not resolve significant effects in our hippocampal slice experiments. Only voxels where static or dynamic modulation were above this threshold are mapped in figure 3C. The maximum of the colorbars in figure 3C was selected as the minimum static or dynamic modulation that could resolve significant effects in our experimental data with 1 kHz carrier.

